# A FZD4/LRP5 agonist restores pericyte coverage and vascular integrity by increasing PDGFB signaling

**DOI:** 10.64898/2026.03.13.711629

**Authors:** Jacklyn Levey, Miranda Howe, Kaia Douglas, Emmanuel Odame, Neal Rajvansh, Ha-Neul Jo, Lingling Zhang, Christina Chung, Heidi Roehrich, Somasekar Seshagiri, Stephane Angers, Zhe Chen, Harald J. Junge

**Affiliations:** Department of Ophthalmology and Visual Neuroscience, University of Minnesota Medical School, Twin Cities, Minneapolis, MN, United States; University of Minnesota, Graduate Program in Molecular, Cellular, and Developmental Biology, and Genetics, Minneapolis, MN; Eppley Institute, University of Nebraska Medical Center, Omaha, NE; University of Wisconsin-Madison, Undergraduate Department of Neurobiology, Madison, WI, USA; University of Colorado Anschutz Medical Campus, Department of Pharmacology, Aurora, CO; Department of Biology, StThomas University, StPaul, MN, USA; Chicago College of Osteopathic Medicine, Chicago, IL, USA; AntlerA Therapeutics, Foster City, CA, USA; Department of Biochemistry, University of Toronto, Toronto, ON, Canada; Terrence Donnelly Centre for Cellular and Biomolecular Research, Toronto, ON, Canada; Leslie Dan Faculty of Pharmacy, University of Toronto, Toronto, ON, Canada; Department of Neuroscience, University of Minnesota, Minneapolis, MN, United States

## Abstract

Pericytes, specialized mural cells of capillaries, fulfill crucial physiological functions including promoting endothelial barrier function and regulating angiogenesis. Pericyte loss or dysfunction represents a central pathological feature in diabetic retinopathy (DR) and is increasingly recognized in neurodegenerative diseases as well as in poor stroke outcomes, underscoring an urgent need for therapies that restore pericyte function or promote their regeneration. Here, we utilized a Frizzled4 (FZD4) and Low-Density Lipoprotein Receptor–Related Protein 5 (LRP5) agonist antibody (F4L5.13) to investigate the functional consequences of mimicking β-catenin-dependent signaling in CNS endothelial cells (ECs), which is physiologically induced by Norrin or WNT7A/B. In platelet-derived growth factor subunit B (*Pdgfb)* EC-specific knockout (ECKO) mice, a model of severe developmental pericyte deficiency with secondary blood-retina barrier (BRB) defects and hemorrhages, F4L5.13 significantly promoted retinal pericyte/mural cell proliferation and coverage, improved BRB function, reduced hemorrhages, and normalized vascular morphology. F4L5.13 restored *Pdgfb* mRNA expression levels from non-recombined cells in *Pdgfb* ECKO retinas. These findings highlight interactions of β-catenin-dependent signaling and PDGFB production, identify a key pharmacodynamic action of F4L5.13 distinct from anti-VEGF therapies, and suggest that FZD4/LRP5 agonists may have uses as a regenerative pharmacology approach that promotes pericyte coverage in the neurovascular unit.

## Introduction

DR is a leading cause of vision impairment in working-age adults (1) and is prominently associated with pericyte loss (2). Pericyte loss or dysfunction is also implicated in glaucoma (3) and has been described in the aging choriocapillaris, linking pericyte loss to dry age-related macular degeneration (4). A reduction or dysfunction of pericytes is furthermore implicated in neurodegenerative disease, small vessel disease, and poor stroke outcomes (5). Together, there is a significant unmet medical need to promote pericyte function, prevent their loss, or regenerate pericyte numbers.

Mural cells are vascular support cells in the blood vessel wall: Vascular smooth muscle cells (VSMCs) cover large vessels (predominantly arteries), transitional mural cells (aka ensheathing pericytes) reside on precapillary arterioles, and pericytes cover capillaries. Loss of pericytes in development causes severe retinal vascular morphogenesis defects, hemorrhages, and BRB defects, reminiscent of DR (6). However, consequences of reduced PC coverage in the adult CNS are largely restricted to mild barrier dysfunction and manifest over long time periods (7). Therefore, it is thought that a reduction of pericyte coverage predisposes the retinal endothelium to inflammatory and pro-angiogenic signals (8, 9), and loss of ensheathing pericytes alters perfusion (10).

Platelet-derived growth factor B (PDGFB), secreted by ECs, is a homodimer composed of two PDGF-B chains and serves as the primary ligand for platelet-derived growth factor receptor-β (PDGFRβ), a receptor tyrosine kinase expressed on mural cells. The binding of PDGFB to PDGFRβ initiates multiple downstream signaling pathways that are essential for mural cell migration, proliferation, and survival (11). PDGFB binds heparan sulfate proteoglycans in the vascular extracellular matrix, a feature essential for localized signaling and precise pericyte recruitment (12, 13). Indiscriminate activation of PDGFB signaling in the retina could have deleterious effects, including disrupting retinal organization (14). AAV-based therapies could deliver PDGFB in a spatially restricted manner but are temporally poorly controlled. Therefore, therapeutic approaches that preserve the spatially restricted nature of PDGFB signaling and are pharmacologically controllable may be required to safely promote mural cell recruitment and proliferation. Approaches that not only prevent further pericyte loss but also stimulate pericyte proliferation could be highly beneficial, as they would regenerate a key cellular component of the neurovascular unit.

The inner BRB is essential for maintaining retinal homeostasis and ensuring the tightly regulated microenvironment required for visual function. This barrier provides selective permeability, tightly controlling the exchange of ions, water, nutrients, hormones, and metabolic waste products between the neural retina and the bloodstream. Structurally, the inner BRB is formed by specialized retinal vascular endothelial cells, which are intercellularly sealed by tight and adherens junction complexes, and supported by pericytes embedded within the basement membrane. Together, these cellular and extracellular components generate a highly restrictive barrier that limits paracellular flux and unspecific transcytosis, while permitting the selective transcellular transport necessary for neuronal survival and function (15). Disruption of the inner BRB results in increased vascular permeability, edema, neuroinflammation, and reduced ERG b-wave (16, 17), and such defects are a hallmark of DR. Endothelial BRB dysfunction is a main cause for diabetic macular edema. Therapeutic strategies that promote endothelial barrier function may be beneficial in the treatment of diabetic retinopathy.

Canonical (i.e., β-catenin-dependent) Norrin and WNT7A/B signaling in vascular ECs are required for CNS angiogenesis and endothelial blood-CNS barrier function (18). In the retina, Norrin is the predominant ligand that induces β-catenin-dependent signaling in ECs via FZD4 (19–21) and requires the tetraspanin TSPAN12 as co-receptor (22–24). Transcriptomic studies show that β-catenin-dependent signaling broadly controls gene expression and EC lineage specialization in accordance with BRB function (25, 26). *Tspan12* gene ablation in mature ECs reveals the important role of Norrin/Frizzled4 signaling in BRB maintenance. Loss of Norrin/Frizzled4 signaling causes BRB defects so severe that cystoid edema is observed in the mouse retina even in the absence of a macula (16, 17, 27). *Fzd4* and *Tspan12* conditional gene ablation studies revealed poorly understood roles of Norrin/Frizzled4 signaling in modulating mural cell coverage in the retina (16, 28).

Recently, we reported that a FZD4/LRP5 agonist antibody modality, F4L5.13, efficiently activates β-catenin-dependent signaling in ECs both *in vivo* and *in vitro* (25, 29). These studies demonstrate the ability of F4L5.13 to restore BRB function in developing and mature endothelial cells. Consistent with restoring BRB function, F4L5.13 achieved complete resolution of cystoid edema in mouse models (17). The pharmacodynamic actions of the emerging class of FZD4/LRP5 and FZD4/LRP6 agonists (30–33) are not well understood. For example, whether FZD4/LRP5 agonists can restore BRB dysfunction caused specifically by reduced pericyte coverage is not known.

Unexpectedly, we find here that F4L5.13 not only alleviated BRB dysfunction associated with mural cell loss but also promoted pericyte/mural cell coverage and reduced hemorrhages. Mechanistically, we found that administration of F4L5.13 boosted *Pdgfb* expression in non-recombined ECs. Restoration of mural cell coverage was at least in part due to increased proliferation of mural cells, suggesting a regenerative aspect of FZD4/LRP5 agonism in the neurovascular unit. Our study highlights interactions of the Norrin/Frizzled4 signaling pathway and PDGFB signaling, and that the pharmacodynamic actions of FZD4/LRP5 agonists are distinct from anti-VEGF therapies and appear highly suitable for the treatment of DR.

## Results

### F4L5.13 restores mural cell coverage in postnatal *Tspan12*^−/−^ mice

In mice, the formation of the three-layered retinal vasculature begins after birth with the development of the superficial vascular plexus from P0-P8, followed by the deep plexus (P7-P12), and the intermediate plexus (P10-P15). In *Tspan12*^−/−^ mice, which are deficient in Norrin/Frizzled4 signaling, the deep vascular plexus fails to form (22) and EC lineage specialization in accordance with BRB induction fails, these phenotypes can be rescued by systemic administration of the FZD4/LRP5 agonist F4L5.13, as the antibody modality does not require the Norrin co-receptor TSPAN12 for signal initiation (29) (17, 25). In addition to angiogenesis defects, Norrin/Frizzled4 signaling is required for proper mural cell coverage (16, 28). To determine if FZD4/LRP5 agonism restored mural cell coverage in *Tspan12*^−/−^ retinas, we administered F4L5.13, i.p., at P4, P7, and P10 (Figure 1A), i.e., during the time window of mural cell recruitment (34). As expected, (17, 25, 29), F4L5.13 significantly restored the formation of the deep vascular plexus and prevented the occurrence of glomeruloid vascular malformations (Figure 1, B-D). While mural cell coverage was continuous in control retinas, in *Tspan12*^−/−^ retinas, the coverage with NG2^+^ (aka CSPG4) mural cells was irregular, patchy, and disconnected; however, glomeruloid vascular malformations retained dense mural cell coverage (Figure 1C). Quantification of mural cell coverage (the area positive for the mural cell marker NG2 divided by the area stained positive with IB4) in the superficial vascular plexus showed significant restoration in animals treated with F4L5.13 (Figure 1, C and E). Increased mural cell coverage was confirmed with RT-qPCR for *Ng2* mRNA (Figure 1F). Similar results were obtained in a cohort of *Tspan12*^−/−^ mice treated with F4L5.13 from P6 to P21 (Supplemental Figure 1A). In this cohort, vascular density in the superficial vascular plexus was more completely restored due to the extended time the animals were treated (Supplemental Figure 1, B-D). Mural cell coverage and *Ng2* mRNA levels were restored to a similar extent in the cohort treated until P12 or until P21. (Supplemental Figure 1E). Together, these findings highlight that increased mural cell coverage represents a poorly understood pharmacodynamic action of the emerging class of FZD4/LRP5 agonists.

**Figure 1.**
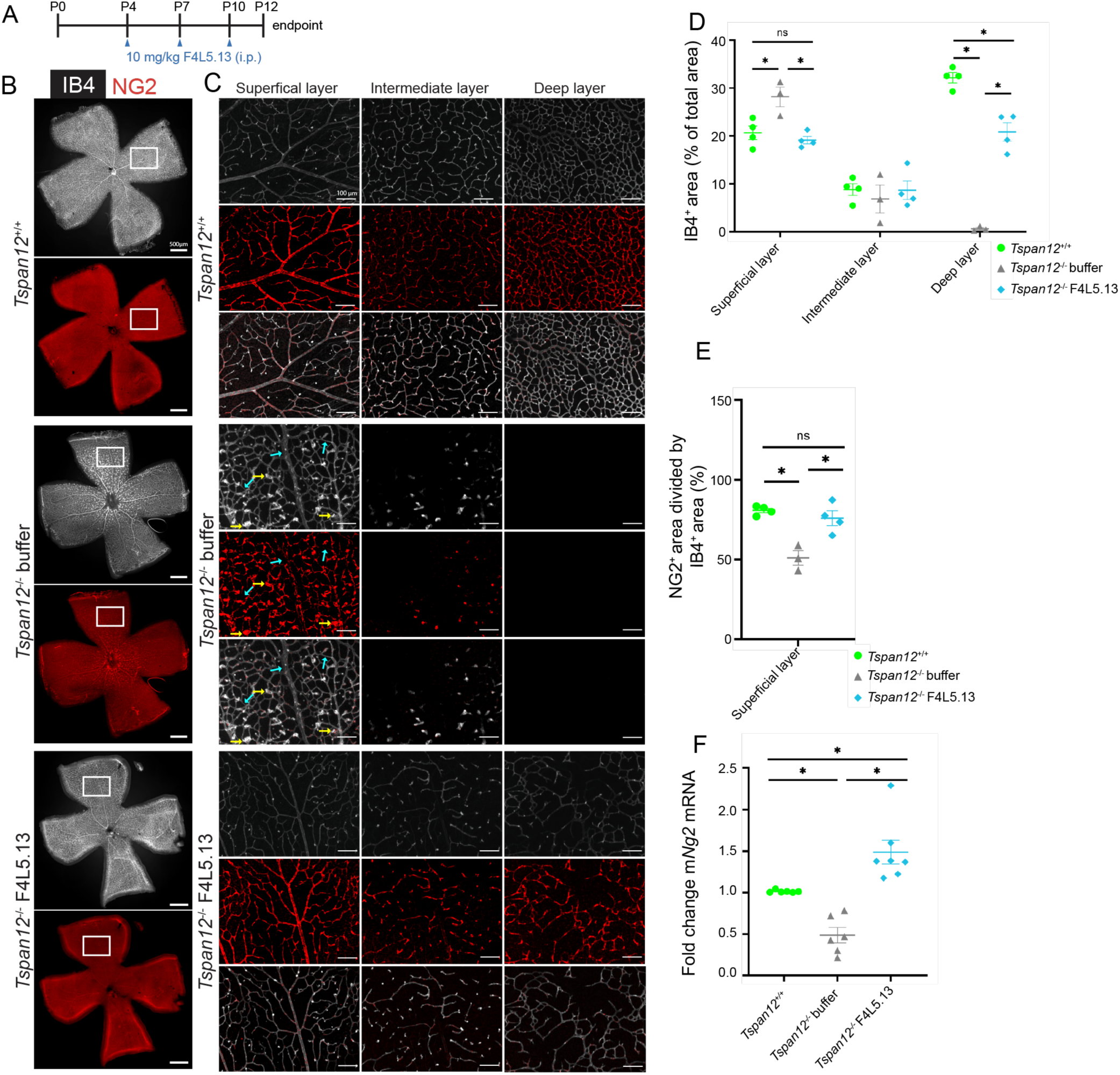
F4L5.13 restores mural cell coverage in *Tspan12*^−/−^ mice. **(A)** Schematic representation of F4L5.13 administration schedule. **(B)** Stitched images of flat-mount retinas obtained from P12 mice of the indicated genotype injected with vehicle or F4L5.13. IB4 (far red channel, grey scale) was used to stain endothelial cells, and anti-NG2 (red) was used to stain mural cells. Boxes outline the area where the 20x image stacks shown in (C) were taken. Scale bar 500 µm. **(C)** 3-D image stacks processed into three separate projections representing the superficial, intermediate, and deep layers of the retinal vasculature. Separate and merged channels (lower panel) are shown. Yellow arrows point to glomeruloid vascular malformations. Cyan arrows point to areas with low mural cell coverage. Scale bar 100 µm. **(D)** Quantification of IB4^+^ area (vascular area) in percent of total retinal area. Four images per retina were averaged, n=3-4 retinas from 3-4 mice per group. Average +/− SE is shown. *P < 0.05 by 1-way ANOVA with Tukey’s post hoc test. **(E)** Quantification of NG2^+^ area divided by IB4^+^ area (to measure mural cell coverage) in each layer of the retinal vasculature. Four images per retina were averaged, n=3-4 retinas from 3-4 mice per group. Average +/− SE is shown. *P < 0.05 by 1-way ANOVA with Tukey’s post hoc test. **(F)** RT-qPCR for *Ng2* from total retinal RNA was normalized to *Gapdh*, n=6-7 retinas from 6-7 mice per group. Average +/− SE is shown. *P < 0.05 by 1-way ANOVA with Tukey’s post hoc test.

### F4L5.13 restores pericyte coverage in *Pdgfb* ECKO retinas

To better understand the effect of FZD4/LRP5 agonists on mural cells, we turned to a disease model in which mural cell loss (predominantly pericytes) is the disease driver. EC-specific *Pdgfb*^fl/fl^ (6) ablation with the Tg(Cdh5-CreERT2)1Rha Cre driver (35) produced *Pdgfb* ECKO mice, i.e., *Pdgfb*^fl/fl^; Cdh5-CreERT2^+^ mice. Cre-negative littermates were used as controls. In *Pdgfb* ECKO mice, loss of PDGFB signaling causes pericyte reduction as a primary defect, which drives abnormal vascular morphogenesis (36), hemorrhages, and barrier dysfunction (9). As expected (6), *Pdgfb* ECKO (tamoxifen at P4-P6, Figure 2A) displayed a significant but variable loss in mural cell coverage (predominantly pericytes). Strikingly, treatment with F4L5.13 significantly restored pericyte coverage in *Pdgfb* ECKO retinas, although the effect in individual retinas was variable (Figure 2B–D). Furthermore, we found that F4L5.13 improved vascular morphogenesis phenotypes in *Pdgfb* ECKO retinas. In mutant retinas, we observed a profoundly malformed superficial vascular plexus with increased vessel diameter and increased vessel density, as well as severely reduced deep vascular plexus development. Remarkably, administration of F4L5.13 significantly improved vascular density in the superficial and deep vascular plexus, albeit a few severely affected retinas showed relatively poor restoration (Figure 2, B–D).

**Figure 2.**
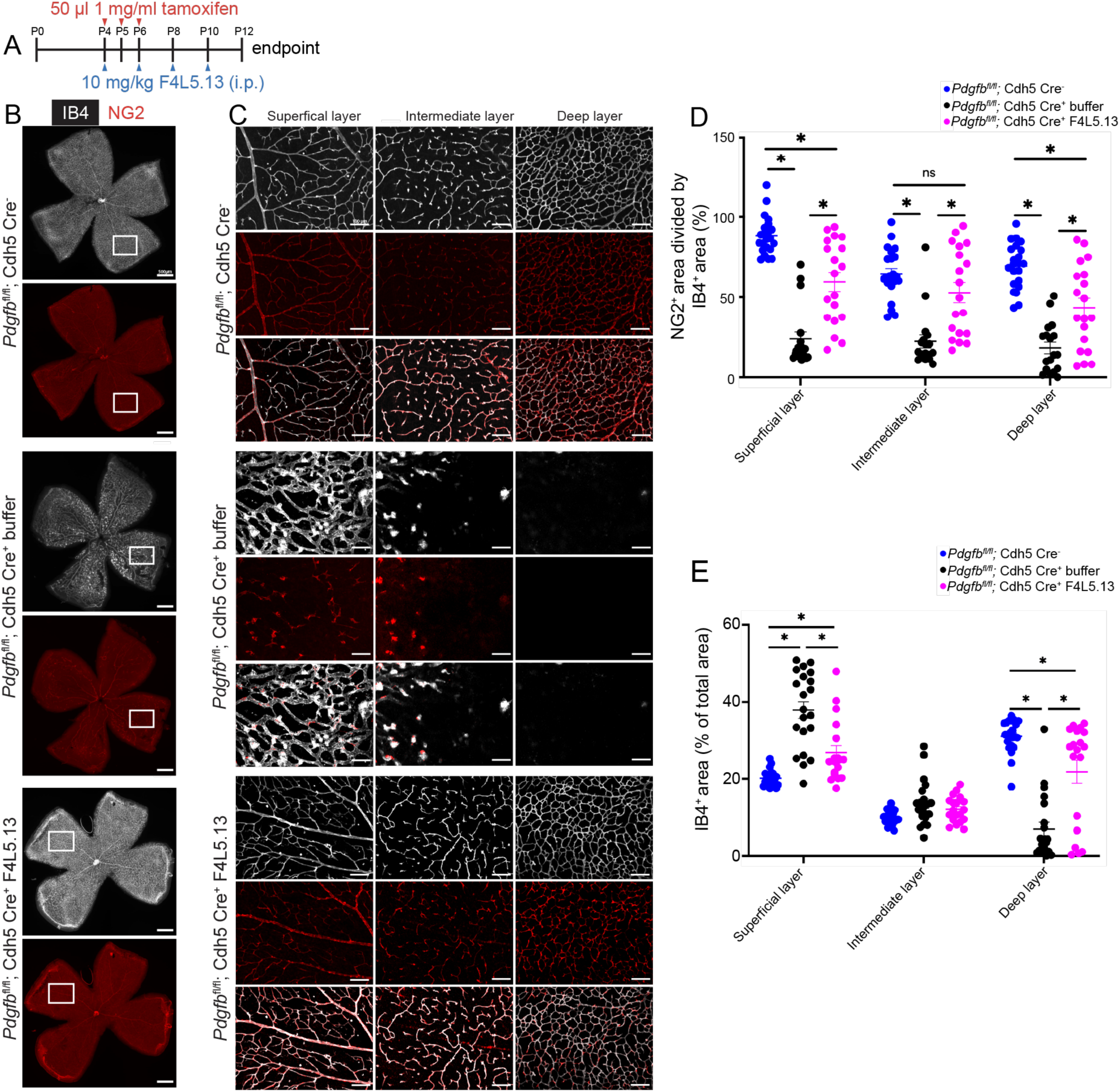
F4L5.13 restores mural cell coverage. **(A)** Schematic representation of F4L5.13 administration schedule. **(B)** Stitched images of flat-mount retinas obtained from P12 mice of the indicated genotype injected with vehicle or F4L5.13. NG2 antibody was used to stain mural cells. Boxes represent the area where the 20x images (C) were taken. Scale bar 500 µm. **(C)** 3-D image stacks processed into three separate projections representing the superficial, intermediate, and deep layers of the retinal vasculature. Separate and merged channels (lower panel) are shown. Scale bar 100 µm. **(D)** Quantification of NG2^+^ area divided by IB4^+^ area (to measure mural cell coverage) in each layer of the retinal vasculature. Four images per retina were averaged, and n=18-22 retinas from 18-22 mice per group were quantified. Average +/− SE is shown. *P < 0.05 by 1-way ANOVA with Tukey’s post hoc test. **(E)** Quantification of IB4^+^ area (vascular area) in percent of total retinal area. Four images per retina were averaged, and n=18-22 retinas from 18-22 mice per group were quantified. *P < 0.05 by 1-way ANOVA with Tukey’s post hoc test.

In addition to reduced pericyte coverage of capillaries, *Pdgfb* ECKO mice exhibited severe abnormalities involving αSMA^+^ mural cells (VSMCs, also myofibroblasts in pathological contexts with fibrosis). In control animals, αSMA^+^ cells were specifically associated with large vessels (predominantly arteries), but in *Pdgfb* ECKO mice αSMA^+^ cells were also associated with retinal capillaries. F4L5.13 treatment restored appropriate localization of αSMA^+^ mural cells to large vessels and reduced coverage with αSMA^+^ mural cells to wild-type levels (Figure 3, A-C). Collectively, these findings suggest that activation of β-catenin-dependent signaling in ECs is a therapeutic strategy to normalize mural cell coverage and distribution.

**Figure 3.**
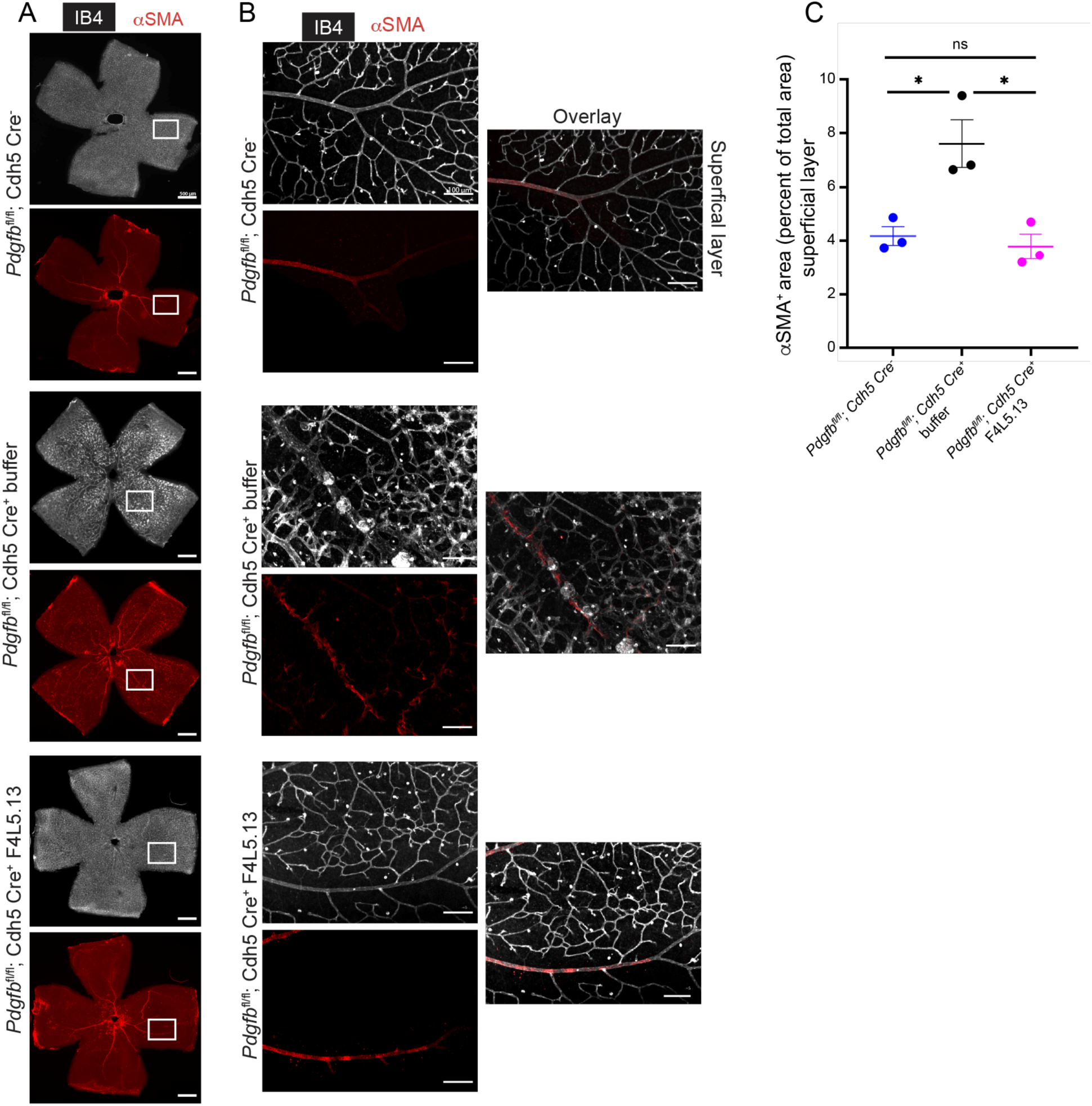
F4L5.13 restores αSMA^+^ cell distribution. **(A)** Stitched images of flat-mount retinas obtained from P12 mice of the indicated genotype injected with vehicle or F4L5.13. Anti-**α**SMA antibody was used to stain VSMCs (also stains myofibroblasts, if present). Boxes outline the area where the 20x images (C) were taken. Scale bar 500 µm. **(B)** A single projection of the superficial vascular plexus was generated from 3-D image stacks. Separate and merged channels (lower panel) are shown. Scale bar 100 µm. **(C)** Quantification of αSMA^+^ area in the superficial vascular plexus in percent of total retinal area. Four images per retina were averaged, and n=3 retinas from 3 mice per group were quantified. Average +/− SE is shown. *P < 0.05 by 1-way ANOVA with Tukey’s post hoc test.

### F4L5.13 restores vascular morphology, integrity, and barrier function in *Pdgfb* ECKO mice

*Pdgfb* ECKO mice display BRB disruption (6, 9). These phenotypes are thought to be independent of a reduction in Norrin/Frizzled4 signaling (see discussion), and indeed, we found that a marker for β-catenin-dependent signaling levels in ECs, CLDN5, was not significantly changed in *Pdgfb* ECKO mice (Supplemental Figure 2, A-C). We assessed BRB function by staining whole-mount retinas for extravasated IgG. *Pdgfb* ECKO mouse retinas displayed a significant increase in IgG extravasation in comparison to controls. BRB function was significantly and consistently improved after F4L5.13 treatment across all animals tested (Figure 4, A-C). To further assess vascular integrity, we examined adherens junctions, focusing on VE-cadherin. In *Pdgfb* ECKO mice, VE-cadherin expression was dysregulated: expression was reduced in larger vessels but elevated in capillaries, and junctional organization was irregular and unorganized. Treatment with F4L5.13 re-established the organized junctional characteristic of healthy vasculature (Supplemental Figure 3, A and B). Together, these findings indicate that F4L5.13 can restore BRB function in pathological contexts that are not due to impaired Norrin/Frizzled4 signaling.

**Figure 4.**
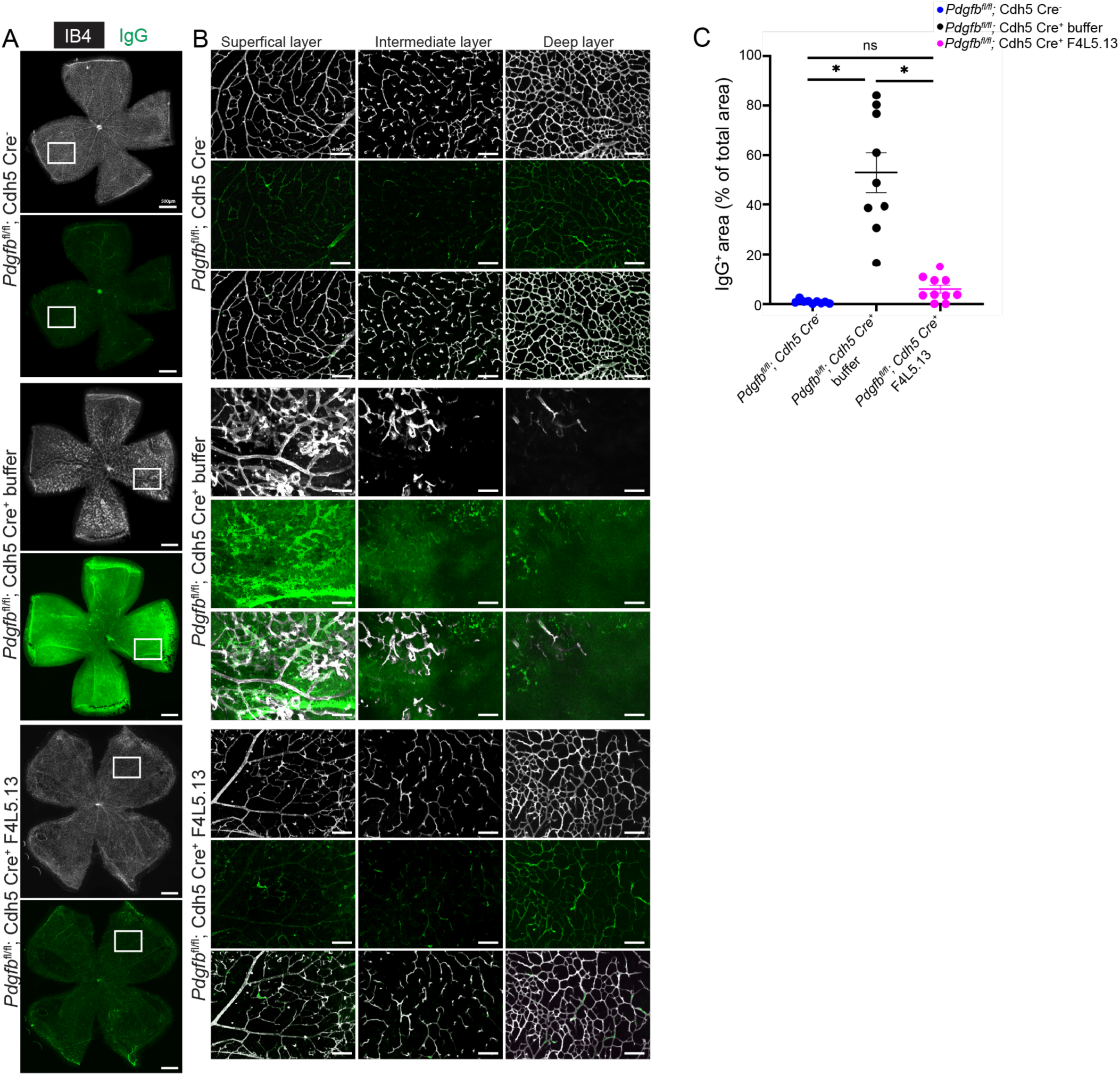
F4L5.13 restores barrier function in *Pdgfb* ECKO mice. **(A)** Stitched images of flat-mount retinas obtained from P12 mice of the indicated genotype injected with vehicle or F4L5.13. Anti-IgG stain was used to assess barrier function. Boxes outline the area where the 20x images shown in (C) were taken. Scale bar 500 µm. **(B)** 3-D image stacks were processed into three separate projections representing the superficial, intermediate, and deep layers of the retinal vasculature. Separate and merged channels (lower panel) are shown. Scale bar 100 µm. **(C)** Quantification of IgG^+^ area in percent of total retinal area. n=9-10 retinas from 9-10 mice per group were quantified. Average +/− SE is shown. *P < 0.05 by 1-way ANOVA with Tukey’s post hoc test.

Loss of pericytes in *Pdgfb* ECKO mice is associated with hemorrhages, providing a model for one of the pathological features of DR. To evaluate if FZD4/LRP5 agonists can alleviate hemorrhages, we stained control retinas, *Pdgfb* ECKO retinas treated with vehicle, and *Pdgfb* ECKO retinas treated with F4L5.13 for the red blood cell (RBC) antigen recognized by the Ter119 antibody. Widespread RBC extravasation was detected in *Pdgfb* ECKO retinas, whereas control retinas showed RBCs within the lumen of blood vessels. Analysis of vascular layers showed that focal spots of leakage were particularly evident in the outer plexiform layer, where sporadic sprouts had begun to form a rudimentary deep vascular plexus. Administration of F4L5.13 substantially reduced RBC extravasation in *Pdgfb* ECKO mice (Figure 5, A-C). This indicates that F4L5.13 not only enhances BRB function but also reinforces overall vascular integrity. This finding is significant, as hemorrhage is a pathological feature of DR and contributes to retinal injury.

**Figure 5.**
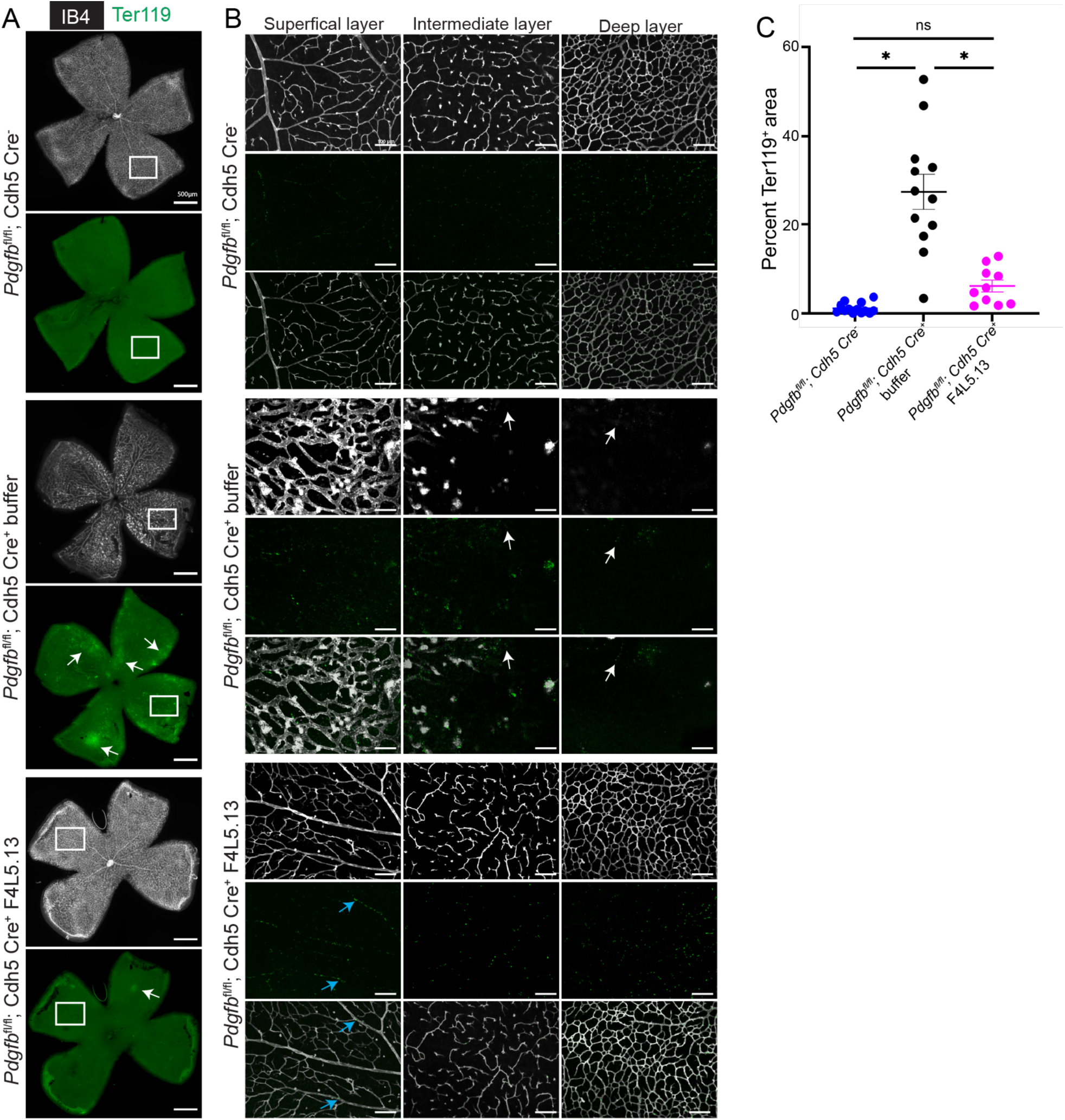
F4L5.13 alleviates hemorrhage in *Pdgfb* ECKO mice. **(A)** Stitched images of flat-mount retinas obtained from P12 mice of the indicated genotype injected with vehicle or F4L5.13. Ter119 antibody was used to stain red blood cells. White arrows mark examples of hemorrhage. Boxes outline the area where the 20x images (C) were taken. Scale bar 500 µm. **(B)** 3-D image stacks processed into three separate projections representing the superficial, intermediate, and deep layers of the retinal vasculature. Separate and merged channels (lower panel) are shown. White arrows mark examples of RBC extravasation, blue arrows mark RBCs inside the lumen of blood vessels. Scale bar 100 µm. **(C)** Quantification of Ter119 signal from 4x stitched images. n=10-13 retinas from 10-13 mice per group were quantified. Average +/− SE is shown. *P < 0.05 by 1-way ANOVA with Tukey’s post hoc test.

### F4L5.13 treatment induces an increase in *Pdgfb* transcription

As *Fzd4* and *Lrp5* are required in ECs to mediate Norrin/Frizzled4 signaling (28, 37), the effects of F4L5.13 on mural cells could be largely indirect via EC transcriptional modulation. To investigate the mechanism by which β-catenin-dependent signaling restores mural cell coverage, we employed a candidate gene approach using qPCR on whole retinal lysates from P12 littermate controls and *Pdgfb* ECKO mice treated with F4L5.13 or vehicle. PDGFB, the key factor in controlling mural cell proliferation, migration, and survival, is expressed in an EC-specific manner in the developing CNS, including the retina (6, 38). RT-qPCR confirmed a mean 62% reduction of *Pdgfb* mRNA in P12 *Pdgfb* ECKO retinas compared with controls and showed that recombination of the *Pdgfb* floxed allele was partial. We used a primer set that anneals within the floxed exon 3, and in exon 4. Therefore, this primer pair is specific for non-recombined *Pdgfb* mRNA. We found that the extent of loss of *Pdgfb* mRNA was variable, matching the variability of pericyte loss in *Pdgfb* ECKO retinas. Importantly, F4L5.13 treatment restored *Pdgfb* mRNA expression in *Pdgfb* ECKO mice (Figure 6A). These results indicate that F4L5.13 boosts *Pdgfb* mRNA expression from non-recombined alleles. To corroborate this finding, we measured levels of *Pdgfb* mRNA in P12 *Tspan12*^−/−^ mice with and without F4L5.13 treatment and compared them with wild type littermate controls. *Pdgfb* mRNA was reduced in *Tspan12*^−/−^ mice and restored in *Tspan12*^−/−^ mice treated with F4L5.13 (Figure 6B). Next, we used a CNS EC-line, bEnd.3 cells, to further substantiate this data. bEnd.3 cells are highly responsive to Norrin or F4L5.13 with respect to upregulating *Axin2* (29), however, these cultured cells do not recapitulate most transcriptional responses to Norrin or F4L5.13 that have been observed by scRNAseq *in vivo* (25). In an RNAseq experiment in a separate study (39), we noticed that bEnd.3 cells express little or no *Lef1,* a factor that mediates transcriptional responses in β-catenin-dependent signaling. To provide sufficient levels of LEF1, we generated a stable bEnd.3 cell population overexpressing LEF1, using a lentiviral vector that introduces LEF1 and resistance to puromycin (Figure 6C). bEnd.3 cells equipped with LEF1 significantly increased *Pdgfb* expression after stimulation with F4L5.13, whereas parental bEnd.3 cells did not respond (Figure 6, D and E). Together, these results indicate that *Pdgfb* is a target (direct or indirect) of β-catenin-dependent signaling in ECs and suggest that FZD4/LRP5 agonists are PDGFB modulators.

**Figure 6.**
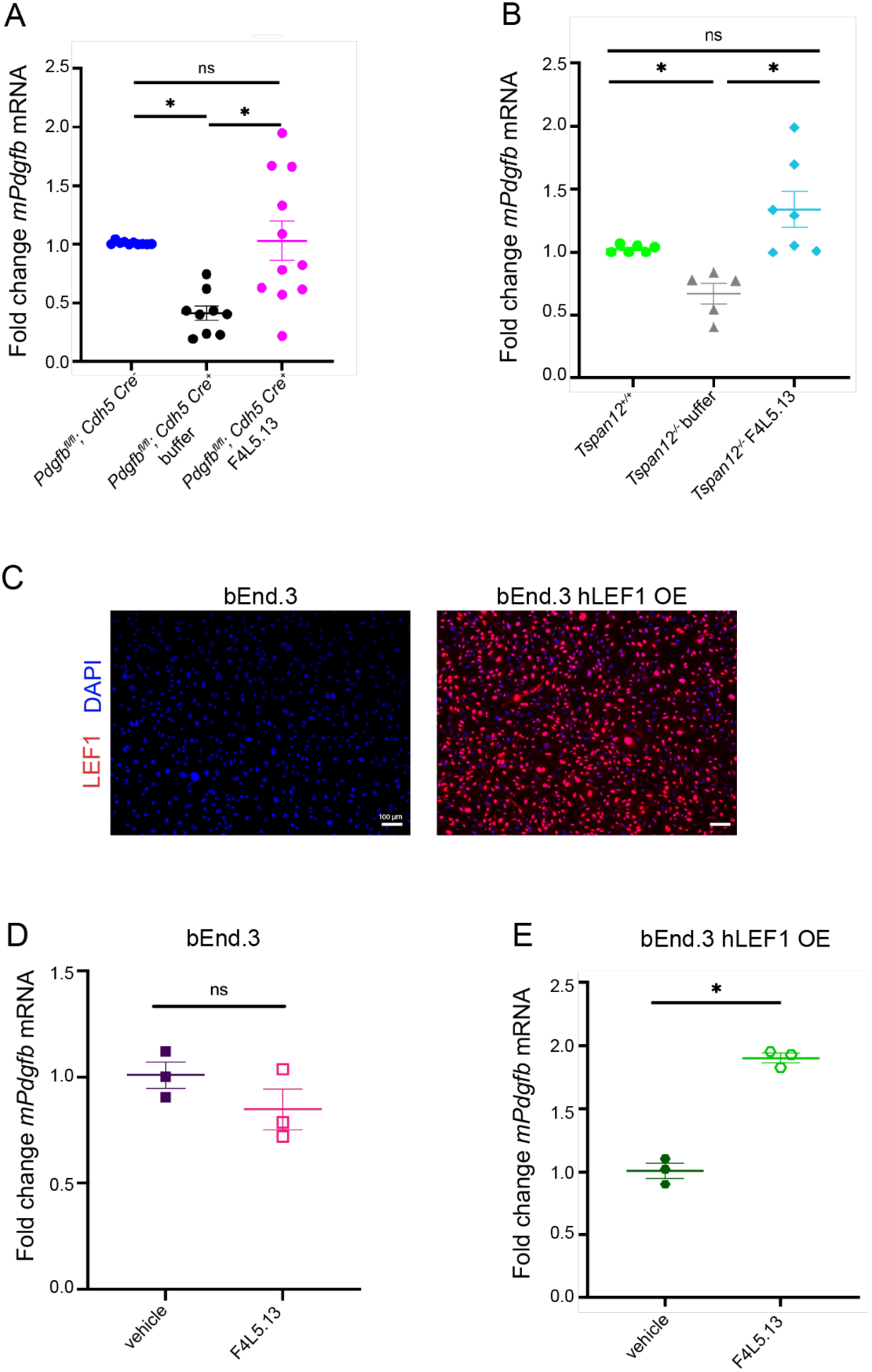
F4L5.13 treatment induces an increase in *Pdgfb* expression. **(A)** Quantification of *Pdgfb* from total retina RNA was normalized to *beta-actin*, n=9-11 retinas from 9-11 mice per group. Average +/− SE is shown. *P < 0.05 by 1-way ANOVA with Tukey’s post hoc test. **(B)** Quantification of *Pdgfb* from total retina RNA was normalized to *gapdh*, n=5-7 retinas from 5-7 mice per group. Average +/− SE is shown. *P < 0.05 by 1-way ANOVA with Tukey’s post hoc test. (**C**) Images of anti-LEF1 stain of parental bEnd.3 cells and a stable population of bEnd.3 cells overexpressing LEF1, generated by lentiviral transduction and selection with puromycin. Scale bar 100 µm. (**D**) Quantification of *Pdgfb* from total RNA of bEnd.3 cells after treatment with vehicle or F4L5.13. 2 technical replicates per biological replicate were averaged, n=3 biological replicates. Average +/− SE is shown. *P < 0.05 by unpaired Student’s t-test. (**E**) Quantification of *Pdgfb* from total RNA of stable bEnd.3-LEF1 cells after treatment with vehicle or F4L5.13. 2 technical replicates per biological replicate were averaged, n=3 biological replicates. Average +/− SE is shown. *P < 0.05 by unpaired Student’s t-test.

### F4L5.13 treatment increases mural cell proliferation

To evaluate how enhanced PDGFB signaling induced by F4L5.13 influences pericyte dynamics in partially recombined *Pdgfb* ECKO retinas, we examined effects on proliferation. PDGFB/PDGFRβ signaling is a major pathway that controls migration and proliferation of mural cells (40, 41). To test the effect of F4L5.13 on mural cell proliferation in control mice and *Pdgfb* ECKO mice treated with vehicle or F4L5.13, three doses of F4L5.13 were administered from P4 to P8, and EdU was administered 4 hours before the P9 endpoint (Figure 7A). EdU positive nuclei of NG2^+^; ERG1^−^ cells at the vascular front and mid-retina were counted separately, as retinal vascular growth follows a central-to-peripheral pattern, with higher proliferation at the angiogenic front than in central regions (Figure 7B). Following F4L5.13 administration, we observed a significant increase in proliferating mural cell numbers (Figure 7C and D), whereas proliferation of mural cells in the mid-retina was not significantly changed (Supplemental Figure 4, A-D). These results suggest that FZD4/LRP5 agonists promote regeneration within the neurovascular unit.

**Figure 7.**
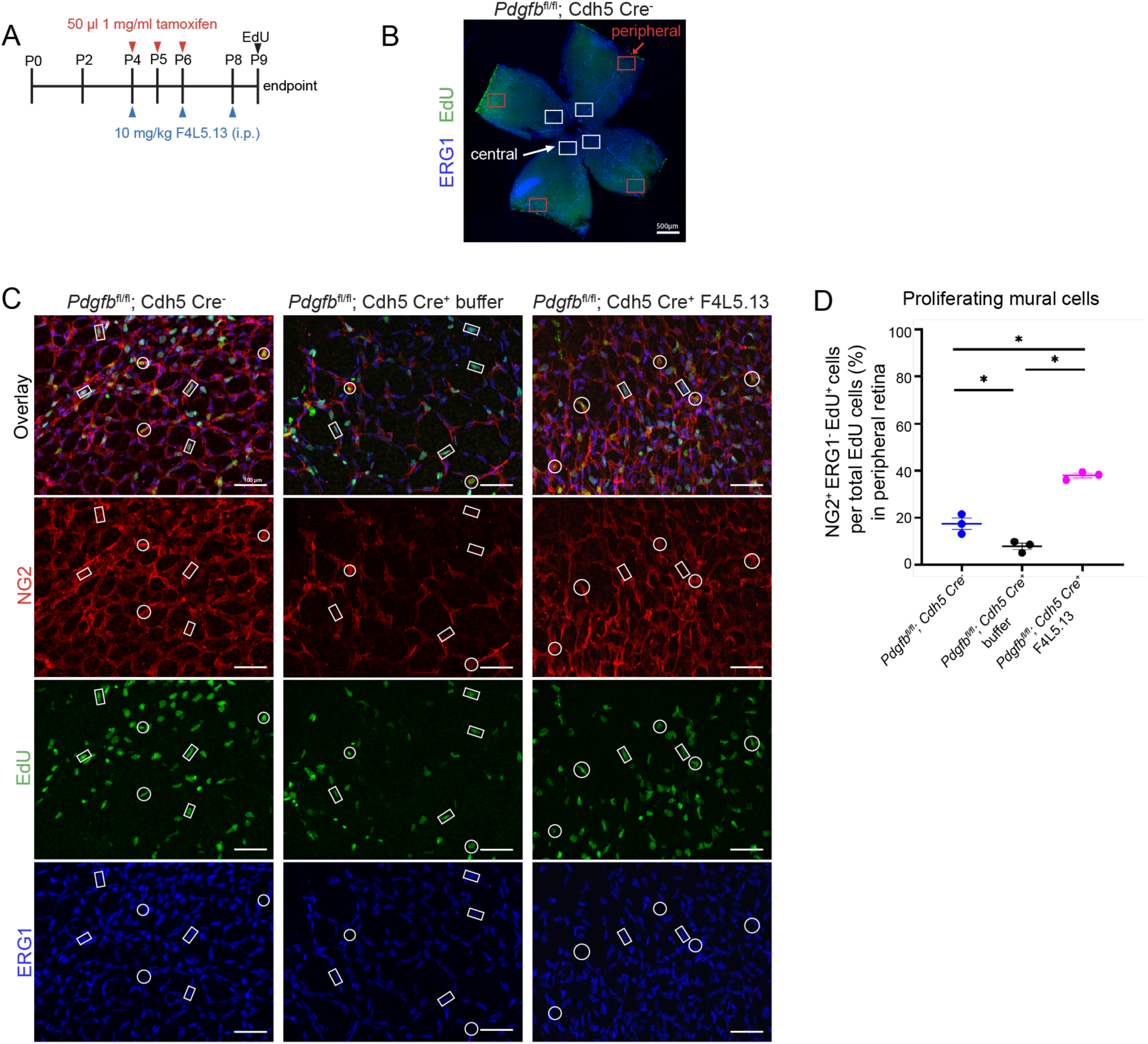
F4L5.13 treatment increases mural cell proliferation. **(A)** Schematic representation of F4L5.13 and EdU administration schedule. EdU was injected 4 hours before harvest. **(B)** Stitched image of flat-mount retina illustrates that proliferation is more prominent in the peripheral retina vs. the central retina. 4 20x images of the peripheral retina were averaged per retina. **(C)** Projections of the superficial vascular plexus in the peripheral retina. Anti-NG2 was used to stain pericytes, anti-ERG1 was used to stain EC nuclei, and EdU was used to identify proliferating cells. White rectangles mark examples for ERG1^+^; EdU^+^ cells. White circles mark examples of NG2^+^; EdU^+^ cells. Scale bar 100 µm. **(D)** Quantification of proliferating mural cells in the peripheral retina. Four images per retina were averaged, and n=3 retinas from 3-mice per group were quantified. Average +/− SE *P < 0.05 by 1-way ANOVA with Tukey’s post hoc test.

## Discussion

DR is a leading cause of impaired vision or blindness in the working age population (1) and is associated with diabetic macular edema (DME) (42) as well as pericyte loss (2). Both non-proliferative and proliferative DR can be accompanied by DME, which is typically caused by endothelial (inner) BRB disruption (43–45). DR is furthermore characterized by hemorrhages, altered vascular morphology (microaneurysms), capillary loss or non-perfusion, lipoprotein extravasation (hard exudates), and ischemia associated with reduced axoplasmic flow in the retinal nerve fiber layer (cotton wool spots) (46). The current standard of care in DME, anti-VEGF therapy, fails to produce adequate therapeutic benefit in a significant subset of patients and does not regenerate pericytes once they are lost. Loss or dysfunction of pericytes is also implicated in stroke, small vessel disease, and neurodegenerative disease (5). There is an unmet medical need for therapies that slow further loss or dysfunction of pericytes or ideally regenerate their number.

In this study, we demonstrate that pharmacological activation of β-catenin-dependent signaling with a FZD4/LRP5 agonist restores retinal mural cell coverage in two genetic models of mural cell loss. The first model, *Tspan12*^−/−^ mice, is characterized by loss of Norrin/Frizzled4 signaling in ECs and moderate mural cell loss on capillaries and large vessels (16). The observation of mural cell coverage defects in *Tspan12*^−/−^mice is consistent with previous reports investigating phenotypes in *Fzd4* mutant mice (28). The second model, *Pdgfb* ECKO mice, is characterized by variable (often severe) mural cell loss, predominantly pericytes. Importantly, the *Pdgfb* ECKO model is thought to be independent of altered β-catenin-dependent signaling in ECs, as expression of the target genes of β-catenin-dependent signaling, CLDN5 (Supplemental Figure 2) and SOX17 (47), are not reduced. Furthermore, scRNAseq data from hypomorphic *Pdgfb*^ret/ret^ mice show a shift towards venous EC lineage specialization without major loss of BBB markers (48), which is not reminiscent of the widespread reduction of BBB markers caused by loss of β-catenin-dependent signaling in ECs (25). The finding that F4L5.13 restores mural cell coverage in a disease model without reduced Norrin/Frizzled4 signaling is significant and implies that FZD4/LRP5 agonists could be therapeutic in disease contexts that are not necessarily caused by loss of β-catenin-dependent signaling in ECs.

FZD4/LRP5 agonists are a novel class of drug candidates (49), whose pharmacodynamic actions are not well understood. Previous studies showed that FZD4/LRP5 agonists promote BRB function (25, 29) and alleviate oxygen-induced retinopathy (29, 50). A FZD4/LRP6 agonist improved stroke outcomes (30). Here, we find that FZD4/LRP5 agonists restore pericytes in two separate genetic disease models. This effect was linked to increased mural cell proliferation (Figure 7), implying a regenerative action of FZD4/LRP5 agonists in the neurovascular unit. Our data suggest that FZD4/LRP5 agonists have the property of being PDGFB modulators. This is significant, as boosting PDGFB expression in ECs leads to retention of PDGFB in the perivascular ECM, avoiding the spatially indiscriminate and likely detrimental (14) activation of retinal cell populations by recombinant PDGFB administration. Even when PDGFB signaling is spatially restricted to vascular cells, the potential for FZD4/LRP5 agonists to elicit adverse fibrotic effects through enhanced PDGFB signaling warrants consideration. While PDGFA is better known in the context of promoting fibrosis and extracellular matrix deposition (51), PDGFB is also implicated in fibrotic remodeling in advanced retinal disease (52). Blood vessel calcification in deep brain structures is linked to PDGFB loss-of-function mutations (53, 54); however, increased circulating PDGFB may also contribute to brain calcification by attenuating receptor signaling through ectodomain shedding (55). The intravitreal administration of FZD4/LRP5 agonists should limit potential adverse effects in the deep brain structures that are, unlike the retinal and choroidal vasculatures, prone to calcification.

The interactions of β-catenin-dependent signaling in ECs, particularly Norrin/Frizzled4 signaling in the retina, and PDGFB signaling, are not well understood. A prior report found a moderate increase of *Pdgfb* and PDGFB in cultured human umbilical vein ECs after activation of β-catenin-dependent signaling, as well as increased mural coverage in glioblastoma (56). In the developing brain, an interaction of retinoic acid signaling, β-catenin-dependent signaling in ECs, and *Pdgfb* was reported (57). Activation of β-catenin-dependent signaling not only alleviates stroke outcomes but is also implicated in mural cell coverage (30, 58). Our observation that a FZD4/LRP5 agonist increases *Pdgfb* mRNA in partially recombined *Pdgfb* ECKO mice as well as in *Tspan12*^−/−^ mice (Figure 6) highlights that β-catenin-dependent signaling in ECs can promote PDGFB signaling in multiple contexts. This provides a mechanistic basis for the pharmacodynamic actions of F4L5.13 on mural cell coverage and mural cell proliferation. The data support a model in which the actions of FZD4/LRP5 agonists on mural cells are indirect through regulation of EC transcripts, of which PDGFB is likely an important mediator, but not necessarily the only one.

Our data shows that reduction of mural cell coverage and loss of *Pdgfb* mRNA are variable in the *Pdgfb* ECKO model (Figure 6) and that F4L5.13 has a significant but variable effect on mural cell restoration (Figure 2). It is likely that animals with the highest degree of recombination efficiency are the most difficult to rescue, as few non-recombined alleles remain whose *Pdgfb* transcription can be boosted by F4L5.13. Interestingly, F4L5.13-mediated rescue of BRB function (IgG staining) and vascular integrity (Ter119 staining) was less variable than its effect on mural cell coverage (Figure 4 and 5). This data implies that not all pharmacodynamic actions of F4L5.13 depend on boosting *Pdgfb* mRNA, and that some of these actions, e.g., on tight or adherens junctions, are independent of mural cell coverage.

Collectively, by simultaneously restoring tight junctions, adherens junctions, and mural cells, FZD4/LRP5 agonists appear to address multiple pathological aspects of BRB dysfunction and reduced vascular integrity with pharmacodynamic actions distinct from anti-VEGF therapies. Whereas anti-VEGF therapies are typically used when VEGF is already elevated and edema or neovascularization is evident, FZD4/LRP5 agonists could potentially have uses at earlier disease stages of DR. In addition, the pharmacodynamic actions of FZD4/LRP5 agonists support potential uses in treating edema in other retinal diseases, including neovascular AMD and retinal vein occlusion.

## Methods

### Sex as a biological variable

Animals of both sexes were used for all studies. The study was not designed or powered to detect sex differences.

### Animals

The *Tspan12* null (Tspan12^tm1.2Hjug^) allele was reported previously and maintained on a C57BL/6J background (22). *Pdgfb* floxed mice (Jackson lab stock 017622) were backcrossed to C57BL/6J. Tg(Cdh5-cre/ERT2)1Rha (38) was used as a Cre driver and was provided by R. Adams under material transfer agreement with https://cancertools.org/. Mice were housed in a specific pathogen-free animal facility.

### Preparation and administration of tamoxifen

Tamoxifen (Sigma- Aldrich, T5648) solution was prepared in sterile corn oil (Sigma-Aldrich, C8267) using a rotator at room temperature overnight, using foil to protect the solution from light. The solution was sterile filtered, aliquoted, and stored frozen at −80°C for no longer than 4 weeks. 50 µl of tamoxifen (1 mg/ml) was injected subcutaneously for 3 days from P4 to P6.

### F4L5.13 antibody administration

F4L5.13 (34) or vehicle (10 mM Histidine, 0.9% sucrose, 150 mM NaCl, pH 6.0) was administered intraperitoneally. *Pdgfb*^fl/fl^; Cdh5-CreERT2^+^, *Tspan12*^−/−^, and BL6J pups received 10 mg/kg at P4, P6, P8 and P10 i.p., as indicated in the respective figures.

### Retinal wholemount immunostaining

Mice were anesthetized using an isoflurane drop jar and euthanized by cervical spine dislocation. Eyes were dissected and mildly fixed in 4% PFA for 15 minutes at room temperature. Retinas were blocked at room temperature with 5% goat serum in PBS with 0.5% Triton X-100 for 2 hours. Staining was performed in blocking buffer at 4°C with shaking overnight using Griffonia Simplicifolia Isolectin B4 Alexa 647 (1:100, Invitrogen, Cat# I32450), anti-NG2 (1:100, Millipore Cat#AB5320), anti-IgG (1:1000, goat anti-mouse Alexa Fluor 488 Cat#A11001), anti-Ter119 (1:200, R&D Cat#MAB1125), anti-CLDN5 Alexa 488 (1:100, Invitrogen Cat#352588), anti-αSMA-Cy3 (1:200, Sigma Cat#C6198), anti-ERG1-Alexa Fluor 647 (1:500, Abcam Cat#ab196149), or anti-CDH5 (1:100, BD biosciences Cat#555289). After overnight incubation, retinas were washed five times for 15 minutes in PBS with 0.1% Triton X-100 while shaking at room temperature. Secondary antibody incubation was performed in blocking buffer at 4°C, shaking overnight using goat anti-rabbit (H+L) Alexa Fluor-555 (1:2000, Invitrogen Cat#A32732), or goat anti-rabbit (H+L) Alexa Fluor 647 (1:2000, Invitrogen Cat#A32733), or goat anti-rat (H+L) Dylight 488 (1:1000, Invitrogen Cat#SA5-10018, or goat anti-rat (H+L) (1:1000, Invitrogen Cat#SA5-10019) Dylight 550. After overnight incubation, retinas were washed five times for 15 minutes in PBS with 0.1% Triton X-100 while shaking at room temperature. Retinas were then postfixed for 15 minutes in 4% PFA at room temperature before mounting. Images were obtained using a Keyence BZ-X810 digital microscope.

### Analysis of vascular leakage and hemorrhage

Using 4x stitched images, a threshold was set based on control littermate staining intensity (Ter119 for hemorrhage, or IgG, for extravasated proteins). The Ter119^+^ or IgG^+^ positive area above threshold was then measured in FIJI to calculate the percent positive area relative to total retinal area.

### Vascular density and mural cell coverage

Vascularized density was calculated from 20x image stacks obtained at 1-μm depth intervals. Multiple images that together represent one of the three vascular layers were projected using the maximum intensity function of ImageJ. A threshold was set for the IB4 intensity using control samples; the same threshold was applied to the experimental genotypes. The IB4-positive area above the threshold was determined and divided by the total area to obtain the percent vascularized area. Mural cell density was calculated in the same manner using 20x stacked images of anti-NG2-stained retinas. For mural cell coverage, the NG2^+^ area was divided by the IB4^+^ area. VSMC density was calculated as SMA^+^ area from 20x stack images as described above. CLDN5 mean gray value was calculated using IB4^+^ area masks. Expression was calculated by applying the vascular mask to the 20x images and measuring the mean gray value within the IB4^+^ area.

### Generation of bEnd.3 hLEF1 overexpressing cells

pLenti-CMV-hLEF1-IRES-puro vector was based on Addgene 103031, deposited by Ghassan Mouneimne (59). hLEF1 was amplified from pcDNA3.3 hLEF1 (23) and inserted into the AscI and NheI sites of the vector. Lentivirus was produced at the University of Minnesota Viral Vector and Cloning Core. 2 mio bEnd.3 cells were seeded in a 10 cm dish and cultured at 5% C02, 37°C, in high glucose DMEM, 10 % FBS, 1 % penicillin/streptomycin. After three days, the cells were infected with 1 ml viral supernatant in 10 ml full medium. After 48 hours, selection with 10 µg/ml puromycin began and was maintained for 1 week with media changes. After that, remaining cells were maintained with 4 µg/ml puromycin. LEF1 was stained in PFA-fixed cells using rabbit anti-LEF1 (Cell Signaling, #2230S 1:100) and goat anti-rabbit Alexa Fluor-555 (Invitrogen, #A21428). Cells were stimulated for 24 hours with 1 µg/ml F4L5.13 or vehicle before extracting total RNA.

### RT-qPCR analysis

Total RNA was purified using RNAzol-RT (ABP Bioscience, FP314) or TRIzol (APB Bioscience, FP312) following the manufacturer’s recommendation. Equal amounts of total RNAs were transcribed into cDNA using the Maxima First Strand cDNA Synthesis Kit for RT (Thermo Fisher Scientific, K-1642), and quantitative PCR was performed using a SYBR green mix. The Δ/Δ^Ct^ method was used for data calculation.

### RT-qPCR primers

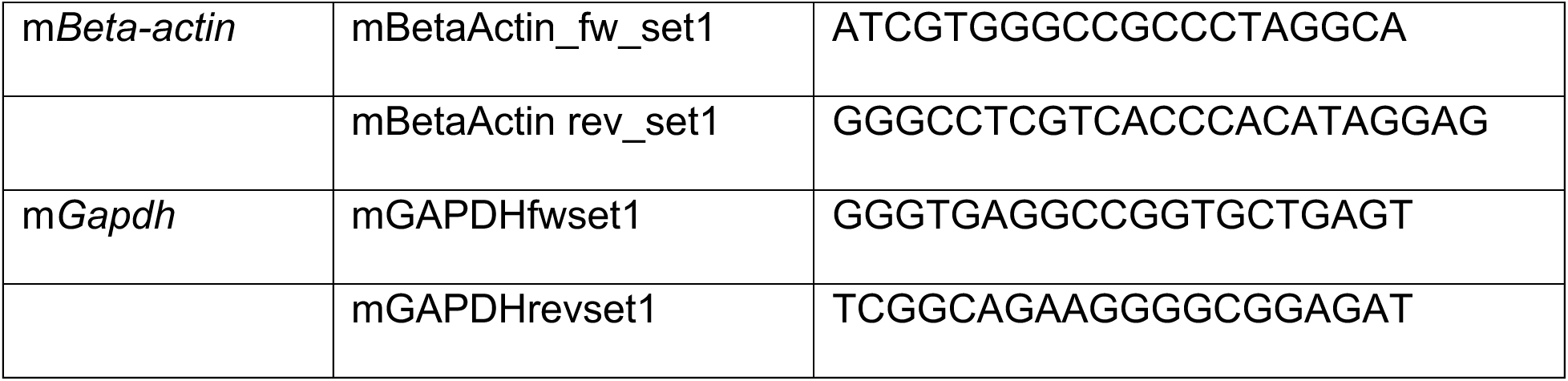
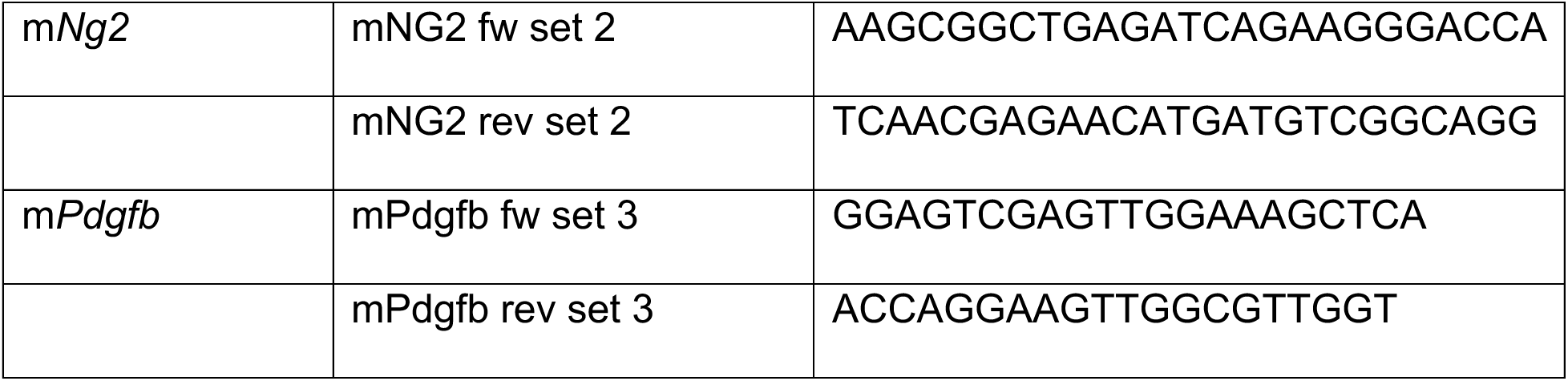

### Analysis of proliferation

In vivo vascular cell proliferation was determined as described (39). In brief, EdU was prepared by dissolving 10 mg of EdU (Invitrogen, A10044) in 100 μl DMSO and 900 μl PBS. Animals were intraperitoneally injected with 10 μl EdU stock solution per gram body weight 4 hours before sacrifice. Retinal wholemount samples were stained with anti-ERG1-647 and anti-NG2 before EdU was detected with the Click-iT-EdU Alexa 488 imaging kit (Invitrogen C10338) according to the manufacturer’s instructions. The total number of EdU^+^ cells was determined. Pericyte proliferation was calculated as percent EdU^+^ cells that were NG2^+^ and ERG1^−^.

### Statistics

A Shapiro-Wilk test was used to assess normality, and Levene’s test was used to evaluate homogeneity of variance. Depending on the distribution and variance, either parametric or non-parametric two-group or multi-group comparisons were performed as described in each figure legend. Parametric tests were homoscedastic or heteroscedastic t-tests and 1-way ANOVA with Tukey’s post hoc analysis. Non-parametric tests were Mann-Whitney U tests and Kruskal-Wallis tests. A p-value <0.05 was considered statistically significant.

### Study approval

All animal protocols were approved by the Animal Care and Use Committee of the University of Minnesota, Twin Cities.

### Data availability

All numeric data supporting the findings of this study are available within the Supporting Data Values files.

## Supporting information

Levey et al., 2026. Supplement

## Author contributions

JL and HJJ wrote the manuscript. JL and HJJ designed the study. JL, MH, KD, EO, NR, HJ, LZ, CC, MH, and ZC conducted experiments. ZC and HR trained personnel or provided the required instrumentation. SS and SA provided the F4L5.13 antibody.

## Competing interests

SS and SA are shareholders of AntlerA Therapeutics. HJJ was a consultant and scientific advisory board member for AntlerA Therapeutics.

## Funding support

This work is the result of NIH funding, in whole or in part, and is subject to the NIH Public Access Policy. Through acceptance of this federal funding, the NIH has been given the right to make the work publicly available in PubMed Central. This study was supported by grants from the NIH (R01EY024261 and R01EY033316 to HJJ and R21DA056728 to ZC), and from the Canadian Institute of Health Research (PJT-175160 to SA).

## References

1. Teo ZL, Tham YC, Yu M, Chee ML, Rim TH, Cheung N, et al. Global Prevalence of Diabetic Retinopathy and Projection of Burden through 2045: Systematic Review and Meta-analysis. Ophthalmology. 2021;128(11):1580–91.

2. Hammes HP, Lin J, Renner O, Shani M, Lundqvist A, Betsholtz C, et al. Pericytes and the pathogenesis of diabetic retinopathy. Diabetes. 2002;51(10):3107–12.

3. Alarcon-Martinez L, Shiga Y, Villafranca-Baughman D, Belforte N, Quintero H, Dotigny F, et al. Pericyte dysfunction and loss of interpericyte tunneling nanotubes promote neurovascular deficits in glaucoma. Proc Natl Acad Sci U S A. 2022;119(7).

4. Nag TC, Gorla S, Kumari C, and Roy TS. Aging of the human choriocapillaris: Evidence that early pericyte damage can trigger endothelial changes. Exp Eye Res. 2021;212:108771.

5. van Splunder H, Villacampa P, Martinez-Romero A, and Graupera M. Pericytes in the disease spotlight. Trends Cell Biol. 2024;34(1):58–71.

6. Enge M, Bjarnegard M, Gerhardt H, Gustafsson E, Kalen M, Asker N, et al. Endothelium-specific platelet-derived growth factor-B ablation mimics diabetic retinopathy. EMBO J. 2002;21(16):4307–16.

7. Vazquez-Liebanas E, Nahar K, Bertuzzi G, Keller A, Betsholtz C, and Mae MA. Adult-induced genetic ablation distinguishes PDGFB roles in blood-brain barrier maintenance and development. J Cereb Blood Flow Metab. 2022;42(2):264–79.

8. Ogura S, Kurata K, Hattori Y, Takase H, Ishiguro-Oonuma T, Hwang Y, et al. Sustained inflammation after pericyte depletion induces irreversible blood-retina barrier breakdown. JCI Insight. 2017;2(3):e90905.

9. Park DY, Lee J, Kim J, Kim K, Hong S, Han S, et al. Plastic roles of pericytes in the blood-retinal barrier. Nat Commun. 2017;8:15296.

10. Hartmann DA, Coelho-Santos V, and Shih AY. Pericyte Control of Blood Flow Across Microvascular Zones in the Central Nervous System. Annu Rev Physiol. 2022;84:331–54.

11. Sweeney MD, Ayyadurai S, and Zlokovic BV. Pericytes of the neurovascular unit: key functions and signaling pathways. Nat Neurosci. 2016;19(6):771–83.

12. Abramsson A, Kurup S, Busse M, Yamada S, Lindblom P, Schallmeiner E, et al. Defective N-sulfation of heparan sulfate proteoglycans limits PDGF-BB binding and pericyte recruitment in vascular development. Genes Dev. 2007;21(3):316–31.

13. Lindblom P, Gerhardt H, Liebner S, Abramsson A, Enge M, Hellstrom M, et al. Endothelial PDGF-B retention is required for proper investment of pericytes in the microvessel wall. Genes Dev. 2003;17(15):1835–40.

14. Edqvist PH, Niklasson M, Vidal-Sanz M, Hallbook F, and Forsberg-Nilsson K. Platelet-derived growth factor over-expression in retinal progenitors results in abnormal retinal vessel formation. PLoS One. 2012;7(8):e42488.

15. Diaz-Coranguez M, Ramos C, and Antonetti DA. The inner blood-retinal barrier: Cellular basis and development. Vision Res. 2017.

16. Zhang C, Lai MB, Pedler MG, Johnson V, Adams RH, Petrash JM, et al. Endothelial Cell-Specific Inactivation of TSPAN12 (Tetraspanin 12) Reveals Pathological Consequences of Barrier Defects in an Otherwise Intact Vasculature. Arterioscler Thromb Vasc Biol. 2018;38(11):2691–705.

17. Zhang L, Levey J, Abedin M, Jo HN, Odame E, Howe M, et al. C1q limits cystoid edema by maintaining basal beta-catenin-dependent signaling and blood-retina barrier function. JCI Insight. 2025.

18. Yemanyi F, Bora K, Blomfield AK, Wang Z, and Chen J. Wnt Signaling in Inner Blood-Retinal Barrier Maintenance. Int J Mol Sci. 2021;22(21).

19. Wang YC, C.; Williams, J.; Zhang, C.; Junge, HJ.; Nathans, J. Interplay of the Norrin and Wnt7a/Wnt7b signaling systems in blood-brain barrier and blood-retina barrier development and maintenance. PNAS In press PMC Journal – In Process. 2018.

20. Xu Q, Wang Y, Dabdoub A, Smallwood PM, Williams J, Woods C, et al. Vascular development in the retina and inner ear: control by Norrin and Frizzled-4, a high-affinity ligand-receptor pair. Cell. 2004;116(6):883–95.

21. Wang Y, Rattner A, Zhou Y, Williams J, Smallwood PM, and Nathans J. Norrin/Frizzled4 signaling in retinal vascular development and blood brain barrier plasticity. Cell. 2012;151(6):1332–44.

22. Junge HJ, Yang S, Burton JB, Paes K, Shu X, French DM, et al. TSPAN12 regulates retinal vascular development by promoting Norrin-but not Wnt-induced FZD4/beta-catenin signaling. Cell. 2009;139(2):299–311.

23. Lai MB, Zhang C, Shi J, Johnson V, Khandan L, McVey J, et al. TSPAN12 Is a Norrin Co-receptor that Amplifies Frizzled4 Ligand Selectivity and Signaling. Cell Rep. 2017;19(13):2809–22.

24. Bruguera ES, Mahoney JP, and Weis WI. The co-receptor Tspan12 directly captures Norrin to promote ligand-specific beta-catenin signaling. bioRxiv. 2024.

25. Zhang L, Abedin M, Jo HN, Levey J, Dinh QC, Chen Z, et al. A Frizzled4-LRP5 agonist promotes blood-retina barrier function by inducing a Norrin-like transcriptional response. iScience. 2023;26(8):107415.

26. Heng JS, Rattner A, Stein-O’Brien GL, Winer BL, Jones BW, Vernon HJ, et al. Hypoxia tolerance in the Norrin-deficient retina and the chronically hypoxic brain studied at single-cell resolution. Proc Natl Acad Sci U S A. 2019;116(18):9103–14.

27. Beck SC, Karlstetter M, Garcia Garrido M, Feng Y, Dannhausen K, Muhlfriedel R, et al. Cystoid edema, neovascularization and inflammatory processes in the murine Norrin-deficient retina. Sci Rep. 2018;8(1):5970.

28. Ye X, Wang Y, Cahill H, Yu M, Badea TC, Smallwood PM, et al. Norrin, frizzled-4, and Lrp5 signaling in endothelial cells controls a genetic program for retinal vascularization. Cell. 2009;139(2):285–98.

29. Chidiac R, Abedin M, Macleod G, Yang A, Thibeault PE, Blazer LL, et al. A Norrin/Wnt surrogate antibody stimulates endothelial cell barrier function and rescues retinopathy. EMBO Mol Med. 2021;13(7):e13977.

30. Ding J, Lee SJ, Vlahos L, Yuki K, Rada CC, van Unen V, et al. Therapeutic blood-brain barrier modulation and stroke treatment by a bioengineered FZD(4)-selective WNT surrogate in mice. Nat Commun. 2023;14(1):2947.

31. Post Y, Lu C, Fletcher RB, Yeh WC, Nguyen H, Lee SJ, et al. Design principles and therapeutic applications of novel synthetic WNT signaling agonists. iScience. 2024;27(6):109938.

32. Nguyen H, Lee SJ, and Li Y. Selective Activation of the Wnt-Signaling Pathway as a Novel Therapy for the Treatment of Diabetic Retinopathy and Other Retinal Vascular Diseases. Pharmaceutics. 2022;14(11).

33. O’Brien S, Chidiac R, and Angers S. Modulation of Wnt-beta-catenin signaling with antibodies: therapeutic opportunities and challenges. Trends Pharmacol Sci. 2023;44(6):354–65.

34. van der Wijk AE, Vogels IMC, van Veen HA, van Noorden CJF, Schlingemann RO, and Klaassen I. Spatial and temporal recruitment of the neurovascular unit during development of the mouse blood-retinal barrier. Tissue Cell. 2018;52:42–50.

35. Wang Y, Nakayama M, Pitulescu ME, Schmidt TS, Bochenek ML, Sakakibara A, et al. Ephrin-B2 controls VEGF-induced angiogenesis and lymphangiogenesis. Nature. 2010;465(7297):483–6.

36. Eilken HM, Dieguez-Hurtado R, Schmidt I, Nakayama M, Jeong HW, Arf H, et al. Pericytes regulate VEGF-induced endothelial sprouting through VEGFR1. Nat Commun. 2017;8(1):1574.

37. Huang W, Li Q, Amiry-Moghaddam M, Hokama M, Sardi SH, Nagao M, et al. Critical Endothelial Regulation by LRP5 during Retinal Vascular Development. PLoS One. 2016;11(3):e0152833.

38. Weng Y, Chen N, Zhang R, He J, Ding X, Cheng G, et al. An integral blood-brain barrier in adulthood relies on microglia-derived PDGFB. Brain Behav Immun. 2024;115:705–17.

39. Levey J, Abedin M, Zhang C, Odame E, Zhang L, Jo HN, et al. The MDM2-p53 axis regulates norrin/frizzled4 signaling and blood-CNS barrier function. Sci Signal. 2025;18(894):eadt0983.

40. Hellstrom M, Kalen M, Lindahl P, Abramsson A, and Betsholtz C. Role of PDGF-B and PDGFR-beta in recruitment of vascular smooth muscle cells and pericytes during embryonic blood vessel formation in the mouse. Development. 1999;126(14):3047–55.

41. Smyth LCD, Highet B, Jansson D, Wu J, Rustenhoven J, Aalderink M, et al. Characterisation of PDGF-BB:PDGFRbeta signalling pathways in human brain pericytes: evidence of disruption in Alzheimer’s disease. Commun Biol. 2022;5(1):235.

42. Yau JW, Rogers SL, Kawasaki R, Lamoureux EL, Kowalski JW, Bek T, et al. Global prevalence and major risk factors of diabetic retinopathy. Diabetes Care. 2012;35(3):556–64.

43. Daruich A, Matet A, Moulin A, Kowalczuk L, Nicolas M, Sellam A, et al. Mechanisms of macular edema: Beyond the surface. Prog Retin Eye Res. 2018;63:20–68.

44. O’Leary F, and Campbell M. The blood-retina barrier in health and disease. FEBS J. 2023;290(4):878–91.

45. Eshaq RS, Aldalati AMZ, Alexander JS, and Harris NR. Diabetic retinopathy: Breaking the barrier. Pathophysiology. 2017;24(4):229–41.

46. Wong TY, Cheung CM, Larsen M, Sharma S, and Simo R. Diabetic retinopathy. Nat Rev Dis Primers. 2016;2:16012.

47. Lin Y, Gahn J, Banerjee K, Dobreva G, Singhal M, Dubrac A, et al. Role of endothelial PDGFB in arterio-venous malformations pathogenesis. Angiogenesis. 2024;27(2):193–209.

48. Mae MA, He L, Nordling S, Vazquez-Liebanas E, Nahar K, Jung B, et al. Single-Cell Analysis of Blood-Brain Barrier Response to Pericyte Loss. Circ Res. 2021;128(4):e46–e62.

49. Wykoff CS, M. AMARONE Shows Promising Outcomes for a Novel Treatment Pathway in DME and nAMD. Retinal Physician. 2024;21(April).

50. Nguyen H, Chen H, Vuppalapaty M, Whisler E, Logas KR, Sampathkumar P, et al. SZN-413, a FZD4 Agonist, as a Potential Novel Therapeutic for the Treatment of Diabetic Retinopathy. Transl Vis Sci Technol. 2022;11(9):19.

51. Klinkhammer BM, Floege J, and Boor P. PDGF in organ fibrosis. Mol Aspects Med. 2018;62:44–62.

52. Liu Y, Noda K, Murata M, Wu D, Kanda A, and Ishida S. Blockade of Platelet-Derived Growth Factor Signaling Inhibits Choroidal Neovascularization and Subretinal Fibrosis in Mice. J Clin Med. 2020;9(7).

53. Keller A, Westenberger A, Sobrido MJ, Garcia-Murias M, Domingo A, Sears RL, et al. Mutations in the gene encoding PDGF-B cause brain calcifications in humans and mice. Nat Genet. 2013;45(9):1077–82.

54. Vanlandewijck M, Lebouvier T, Andaloussi Mae M, Nahar K, Hornemann S, Kenkel D, et al. Functional Characterization of Germline Mutations in PDGFB and PDGFRB in Primary Familial Brain Calcification. PLoS One. 2015;10(11):e0143407.

55. Wang J, Fang CL, Noller K, Wei Z, Liu G, Shen K, et al. Bone-derived PDGF-BB drives brain vascular calcification in male mice. J Clin Invest. 2023;133(23).

56. Reis M, Czupalla CJ, Ziegler N, Devraj K, Zinke J, Seidel S, et al. Endothelial Wnt/beta-catenin signaling inhibits glioma angiogenesis and normalizes tumor blood vessels by inducing PDGF-B expression. J Exp Med. 2012;209(9):1611–27.

57. Bonney S, Dennison BJC, Wendlandt M, and Siegenthaler JA. Retinoic Acid Regulates Endothelial beta-catenin Expression and Pericyte Numbers in the Developing Brain Vasculature. Front Cell Neurosci. 2018;12:476.

58. Chang J, Mancuso MR, Maier C, Liang X, Yuki K, Yang L, et al. Gpr124 is essential for blood-brain barrier integrity in central nervous system disease. Nat Med. 2017;23(4):450–60.

59. Padilla-Rodriguez M, Parker SS, Adams DG, Westerling T, Puleo JI, Watson AW, et al. The actin cytoskeletal architecture of estrogen receptor positive breast cancer cells suppresses invasion. Nat Commun. 2018;9(1):2980.

