## Supplementary material for "A FZD4/LRP5 agonist restores pericyte coverage and vascular integrity by increasing PDGFB signaling": Levey et al., 2026. Supplement

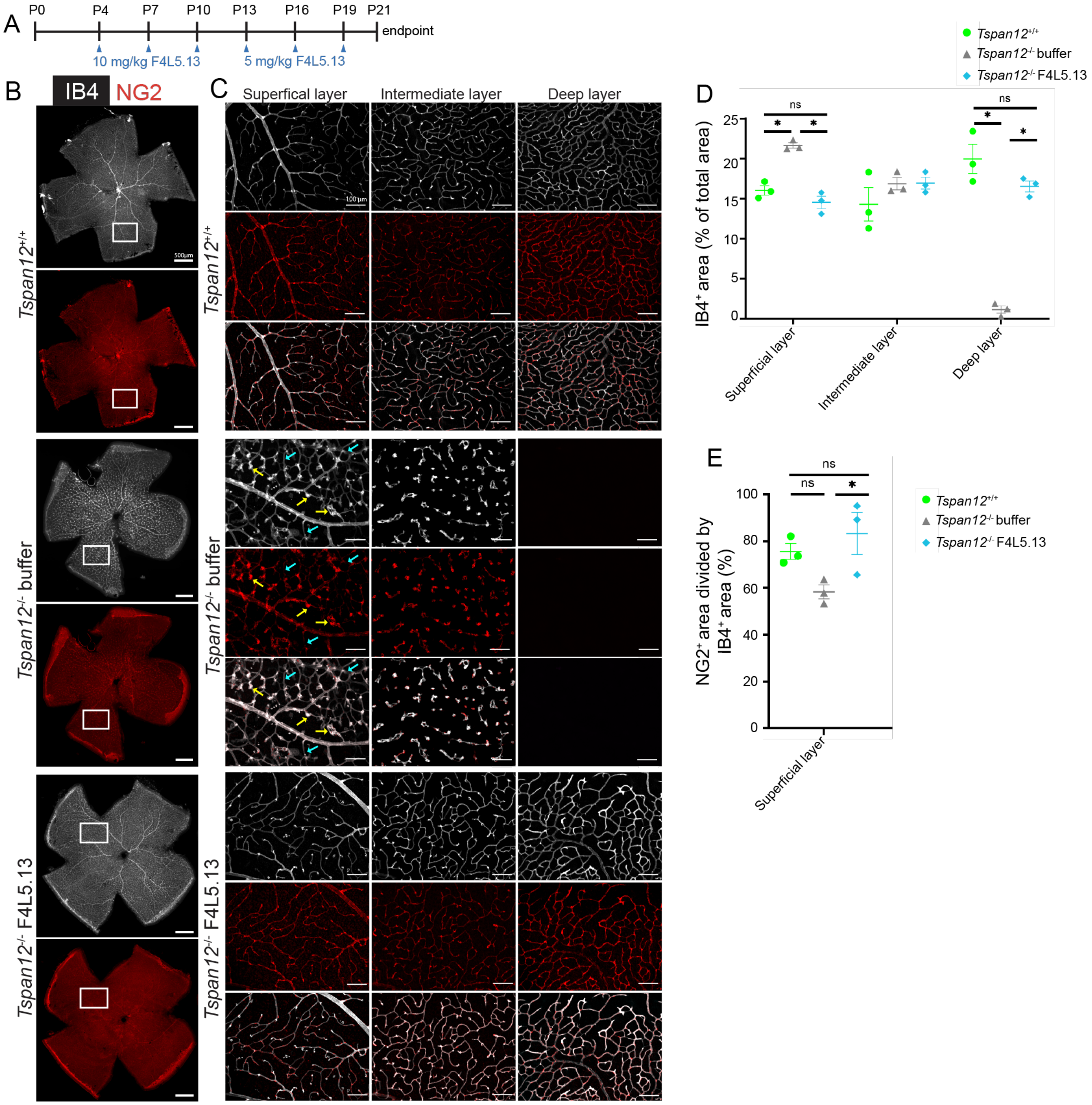


**Supplemental Figure 1. F4L5.13 restores mural cell coverage in P21 *Tspan12*$^{-/-}$ mice.**

**(A)** Schematic representation of F4L5.13 administration schedule. **(B)** Stitched images of flat-mount retinas obtained from P21 mice of the indicated genotype injected with vehicle or F4L5.13. IB4 (far red channel, grey scale) was used to stain endothelial cells, and anti-NG2 (red) was used to stain mural cells. Boxes outline the area where the 20x image stacks show in (C) were taken. Scale bar 500 µm. **(C)** 3-D image stacks processed into three separate projections representing the superficial, intermediate, and deep layers of the retinal vasculature. Separate and merged channels (lower panel) are shown. Yellow arrows point to glomeruloid vascular malformations. Cyan arrows point to areas with low mural cell coverage. Scale bar 100 µm. **(D)** Quantification of IB4$^{+}$ area (vascular area) in percent of total retinal area. Four images per retina were averaged, n=3 retinas from 3 mice per group. Average +/- SE is shown. *$P < 0.05$ by 1-way ANOVA with Tukey's post hoc test. **(E)** Quantification of $NG2^{+}$ area divided by $IB4^{+}$ area (to measure mural cell coverage) in each layer of the retinal vasculature. Four images per retina were averaged, n=3 retinas from 3 mice per group. Average +/- SE is shown. *$P < 0.05$ by 1-way ANOVA with Tukey's post hoc test.

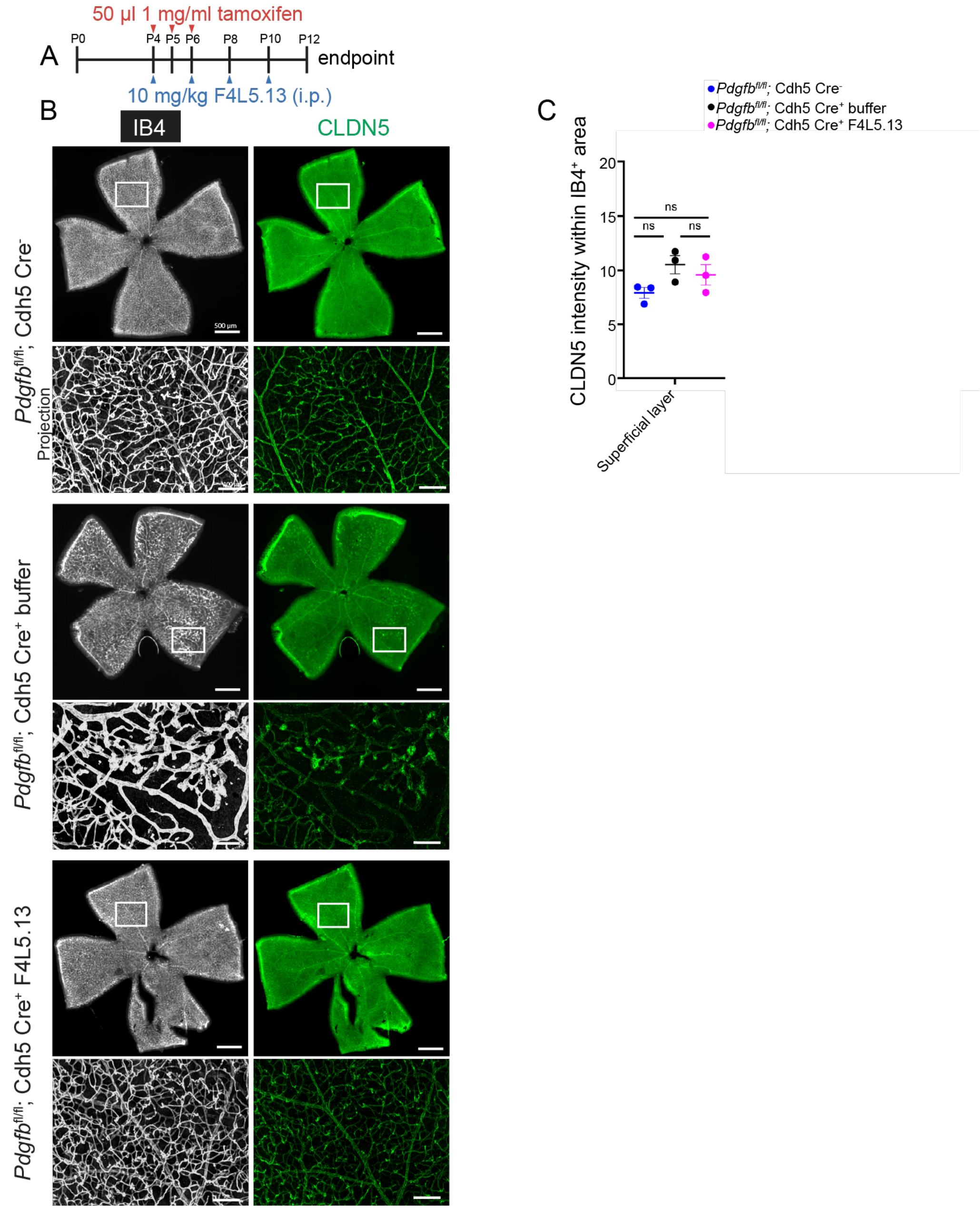


**Supplemental Figure 2. The Wnt signaling marker CLDN5 is not changed in *Pdgfb* ECKO retinas.**

**(A)** Schematic representation of F4L5.13 administration schedule. **(B)** Stitched images of flat-mount retinas obtained from P21 mice of the indicated genotype injected with vehicle or F4L5.13. IB4 (far red channel, grey scale) was used to stain ECs and anti-CLDN5 (red) was used to stain EC tight junctions. Boxes outline examples of areas used for quantification. Scale bar 500 µm. **(C)** Quantification of CLDN5 intensity within the IB4$^+$ area. Four images per retina were averaged, n=3 retinas from 3 mice per group. Average +/- SE is shown. *P < 0.05 by 1-way ANOVA with Tukey's post hoc test.

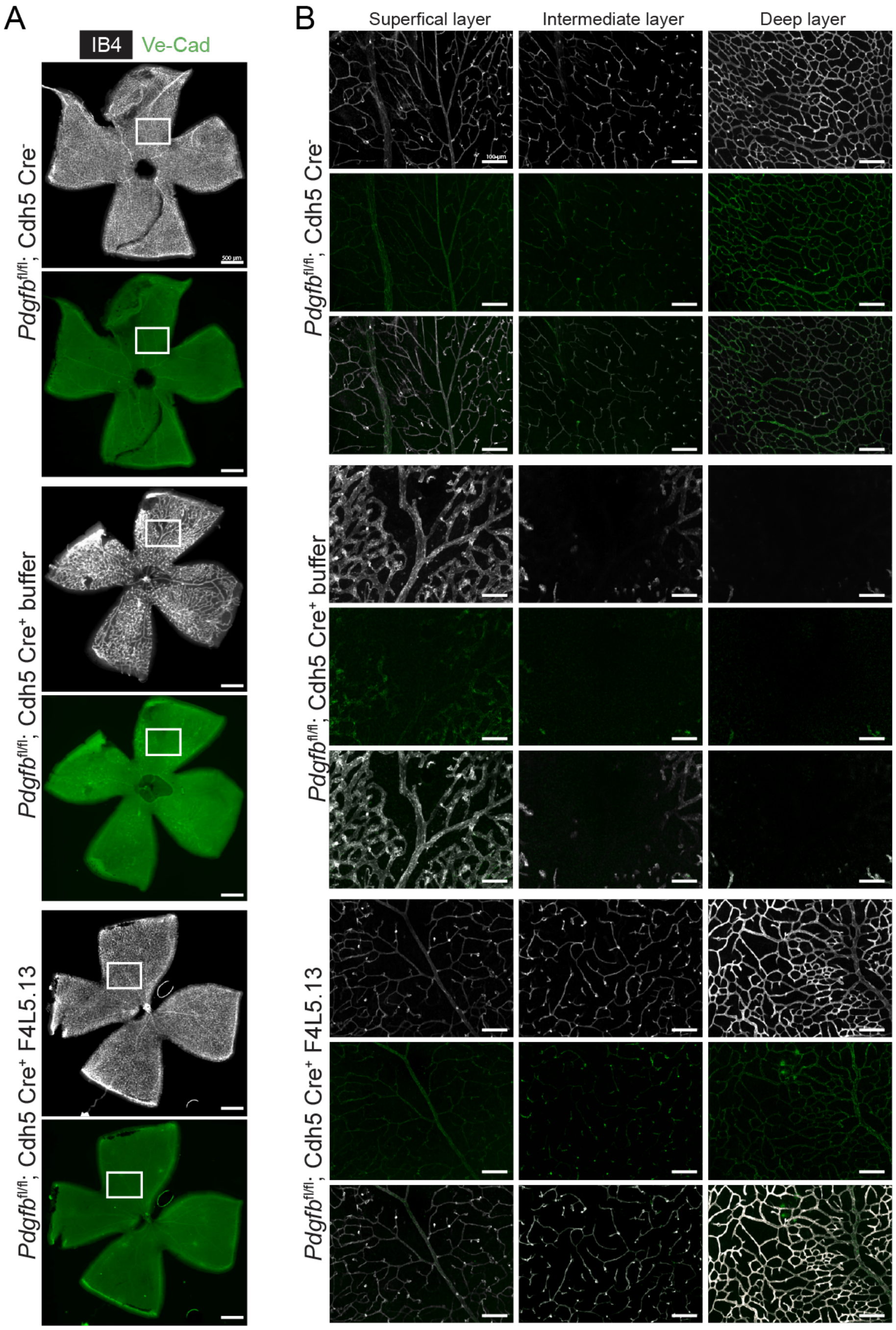


**Supplemental Figure 3. F4L5.13 alleviates adherens junction disorganization.**
**(A)** Stitched images of flat-mount retinas obtained from mice of the indicated genotype injected with vehicle or F4L5.13. IB4 (far red channel, grey scale) was used to stain ECs and anti-CDH5 (green) was used to stain EC adherens junctions. Boxes outline examples of areas shown enlarged in panel B. Scale bar 500 μm. **(B)** 3-D image stacks processed into three separate projections representing the superficial, intermediate, and deep layers of the retinal vasculature. Separate and merged channels (lower panel) are shown. Representative of 3-4 retinas from 3 -4 mice per group with similar results. Scale bar 100 μm.

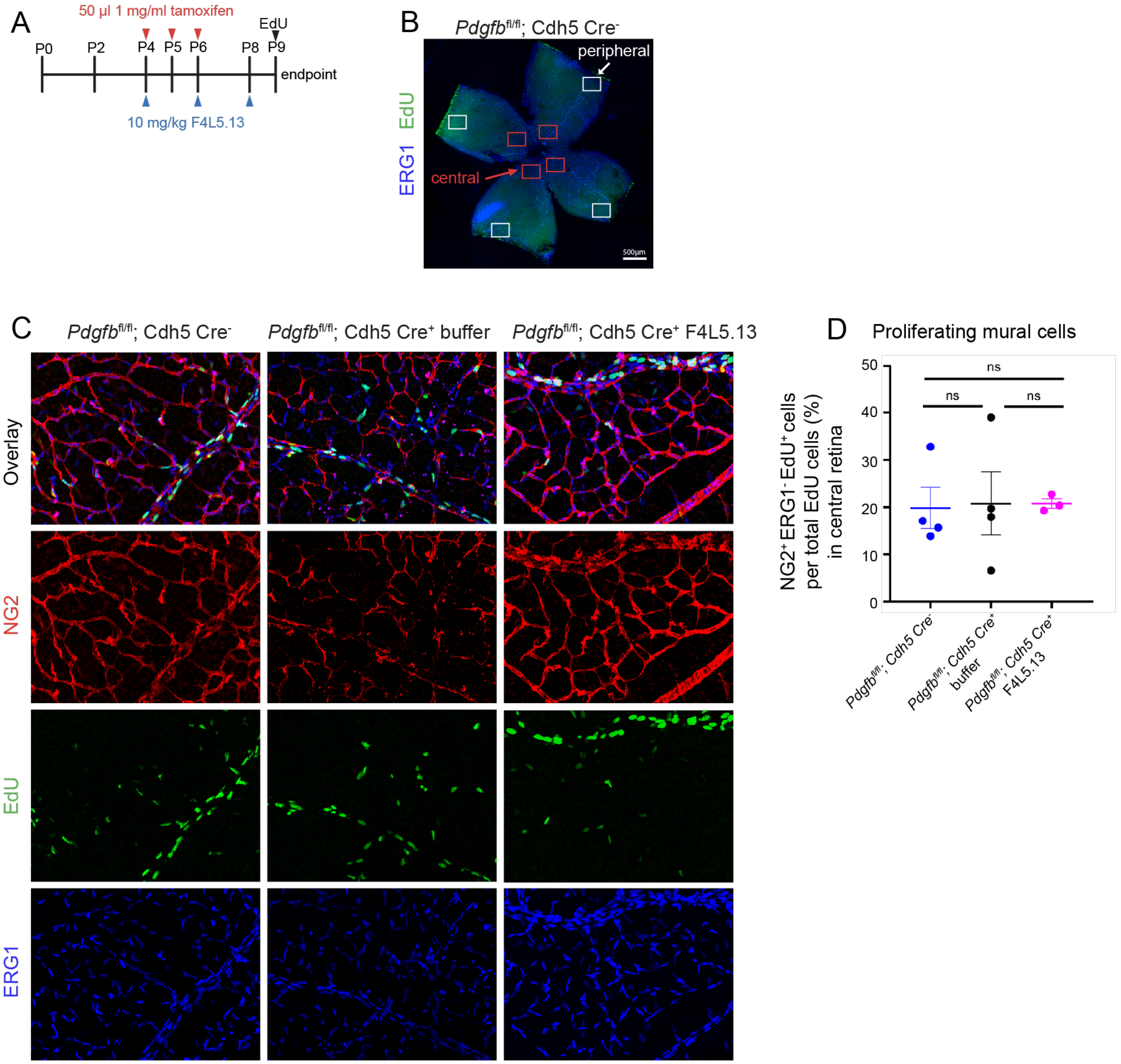


**Supplemental Figure 4. No effect of F4L5.13 treatment on mural cell proliferation in the P9 central retina.**

**(A)** Schematic representation of F4L5.13 and EdU administration schedule. EdU was injected 4 hours before harvest. **(B)** Stitched image of flat-mount retina illustrates that proliferation is more prominent in the peripheral retina vs. the central retina. 4 20x images of the central retina were averaged per retina. **(C)** Projections of the superficial vascular plexus in the peripheral retina. Anti-NG2 was used to stain pericytes, anti-ERG1 was used to stain EC nuclei, and EdU was used to identify proliferating cells. White rectangles mark examples for ERG1$^+$; EdU$^+$ cells. White circles mark examples of NG2$^+$; EdU$^+$ cells. Scale bar 100 µm. **(D)** Quantification of proliferating mural cells in the central retina. Four images per retina were averaged, and n=3-4 retinas from 3-4 mice per group were quantified. Average +/- SE *$P < 0.05$ by 1-way ANOVA with Tukey's post hoc test.
